# Sex difference in hip adduction during the stance phase of running: A swing phase problem?

**DOI:** 10.1101/2020.12.14.422526

**Authors:** Jia Liu, George J Salem, Christopher M Powers

## Abstract

**Purpose:** The purpose of the current study was to: 1) evaluate sex differences in peak hip adduction during the late swing and stance phases of running and 2) determine if peak hip adduction during late swing is predictive of peak hip adduction during stance.

**Methods:** 15 female and 16 male heel strike runners ran over ground at a speed of 4 m/s. Hip joint kinematics during running were quantified using a 3D motion capture system. Sex differences in peak hip adduction during the late swing and stance phases were compared using independent sample t-tests. Linear regression analysis was used to determine the relationship between late swing and stance phase hip adduction.

**Results:** Compared to males, females exhibited significantly greater peak hip adduction during both the late swing (8.5 ± 2.6 vs 6.1 ± 2.8°, p = 0.019) and stance phases of running (13.3 ± 4.2 vs 9.6 ± 3.4°, p = 0.011). Furthermore, late swing peak hip adduction was predictive of subsequent stance phase peak hip adduction (r = 0.63, p < 0.001).

**Conclusion:** Sex differences in hip adduction during stance are influenced in part by late swing phase hip adduction. Further studies are needed to identify potential causes of excessive hip adduction during the late swing phase of running.

## INTRODUCTION

Excessive hip adduction is a common movement impairment that has been reported to underlie various running injuries (1–6). Compared to males, females exhibit greater degrees of hip adduction regardless of speed (7,8), inclination (8), or mode of running (i.e., overground or treadmill) (9). The higher magnitude of hip adduction observed in female runners is thought to contribute to the reported sex differences in the incidence of patellofemoral pain (1,10) and iliotibial band syndrome (3–5).

Most studies that have examined sex differences in frontal plane hip kinematics have focused primarily on the stance phase of running (7,11–16). However, females also have been reported to exhibit higher degrees of hip adduction at initial contact (7,13,16). This suggests that the tendency of female runners to exhibit greater degrees of hip adduction is not just limited to stance, but also may be present during the late swing phase of running.

During the late swing phase of running, hip adduction serves to align the base of support with the center of mass in preparation for ground contact (9). Given that females have distinct differences in pelvis morphology compared to males (i.e., acetabulum orientation and bilateral hip joint width (17–20)), it is possible that greater degrees of hip adduction would be required to position the foot relative to the center of mass to achieve stance-phase stability. An increase in swing phase hip adduction however may bias the hip towards greater degrees of hip adduction during stance.

Diminished hip abductor strength is commonly believed to contribute to excessive hip adduction during the stance phase of running. However, studies that have examined this relationship have reported weak associations (12) or no associations at all (21–24). It is conceivable that stance phase hip adduction, may be influenced to a greater degree by the frontal plane orientation of the hip in late swing phase hip adduction. To further explore this premise, the purpose of the current study was two-fold: 1) to evaluate sex differences in peak hip adduction during the late swing and stance phases of running and 2) determine if peak hip adduction during late swing is predictive of peak hip adduction during stance. Based on previous reports in this area, we hypothesized that, compared to males, females would demonstrate greater peak hip adduction during the stance and late swing phases of running. In addition, we hypothesized that peak hip adduction during the late swing phase of running would be predictive of stance phase values for both males and females. Information gained from this study may provide additional insight into potential causes of sex differences in lower limb kinematics that are related to running injuries.

## METHODS

### Participants

Thirty-one recreational runners participated in this study (16 males and 15 females; TABLE 1). To be eligible for the study, participants must have been between 18-45 years of age, and currently running at least 16 km per week. All participants were natural heel strikers (e.g., runner who contacted the ground with the rear third of their foot first), which was verified using sagittal plane images from high speed video (120 Hz). Only heel strikers were recruited because previous research has shown that hip adduction during running with forefoot striking is smaller than with heel striking (25).

**TABLE 1.** Participant Demographic.

|  | Females<br>(n=15) | Males<br>(n=16) |
| --- | --- | --- |
| Age (years) | 25.2 ± 3.4 | 25.6 ± 4.6 |
| Height (m)* | 1.7 ± 0.1 | 1.8 ± 0.1 |
| Weight (kg)* | 59.2 ± 8.5 | 77.5 ± 9.9 |
| Running distance (km/week) | 25.3 ± 9.2 | 34.4 ± 29.2 |
| Stride Length (m) | 2.34 ± 0.18 | 2.44 ± 0.20 |
| Stride Width (m) | 0.07 ± 0.05 | 0.10 ± 0.05 |

Values are mean ± SD.
\*: *significant sex difference, $p < 0.05$*

Potential participants were excluded if they reported any of the following: (1) current lower extremity or low back pain; (2) previous history of lower extremity surgery, fracture, osteoarthritis, or hip dysplasia, or (3) any lower extremity pathology that caused pain or discomfort during running within 6 months prior to participation. An a priori power analysis using **previous** data obtained from healthy young female and male runners indicated that 11 subjects per group would be sufficient to detect differences and correlations in variables of interest with a statistical power of 80% (using an alpha level of 0.05).

### Instrumentation

Three-dimensional lower extremity kinematics during running were collected using an 11-camera motion capture system (Qualisys, Gothenburg, Sweden) at a sampling rate of 250 Hz. Ground reaction force data were obtained at a rate of 1500 Hz using a single force plate (AMTI, Newton, MA). Kinematic and ground reaction force data were collected and synchronized using a motion capturing software (Qualisys Track Manager version 2.12).

### Procedure

Data were collected at the Jacquelin Perry Musculoskeletal Biomechanics Research Laboratory at the University of Southern California. Prior to data collection, participants were informed as to the objectives, procedures, and potential risks of participation in the study and provided written informed consent as approved by the Health Science Institutional Review Board of the University of Southern California.

Data were obtained from each participant’s dominant leg which was defined as the leg they preferred to use when kicking a ball. Prior to data collection, 21 anatomical markers (reflective 14-mm diameter) were placed on the following bony landmarks: distal phalange of second toes, first and fifth metatarsal heads, medial and lateral malleoli, medial and lateral epicondyles of femurs, greater trochanters, iliac crests, L5-S1 junction, and anterior superior iliac spines (ASISs). In addition, tracking marker clusters mounted on semi-rigid plastic plates were placed on the posterior sacrum and the lateral surfaces of the participant’s thighs, shanks, and heel counters of the shoes. A standing calibration trial was obtained to define the segmental coordinate systems and joint axes. After the calibration trial, anatomical markers were removed, except for those at the ASISs, iliac crests, and L5-S1 junction.

Participants were instructed to run over ground at a controlled speed of 4 m/s along a 14-meter runway. A successful trial was defined when the running speed was within ± 5% of the target speed and the foot of the dominant leg fell within the borders of the force plate. A total of 3 successful trials were collected from each participant.

### Data Analysis

Kinematic data were low pass filtered at 20Hz using a fourth-order Butterworth filter (26). Visual 3D software (C-Motion, Rockville, MD) was used to quantify 3-D hip joint kinematics. using a Cardan rotation sequence of flexion/extension, abduction/adduction, and internal/external rotation (27). Peak hip adduction during the late swing phase of running (i.e., last 1/3 of swing phase) and the deceleration phase of stance (i.e., from heel strike to maximum knee flexion during stance) were identified for each trial. Data obtained from the 3 trials were averaged for statistical analysis.

### Statistical Analysis

Regarding the first hypothesis, sex differences in peak adduction during the late swing phase and stance phase of running were evaluated using independent sample t-tests. To evaluate whether peak hip adduction during late swing was predictive of peak adduction during stance (second hypothesis) multiple linear regression analysis was used. First we tested for the presence of a sex interaction, Since no interaction was present, males and females were combined in the final simple linear regression model. All statistical analyses were performed using R Statistical Software (R Foundation for Statistical Computing, Vienna, Austria), using significance level α=0.05.

## RESULTS

Hip adduction time series data for males and females are presented in **FIGURE 1**. Compared to males, females exhibited significantly greater peak hip adduction during both the late swing phase (8.5 ± 2.6 vs 6.1 ± 2.8°, p = 0.019, **FIGURE 2a**) and stance phase of running (13.3 ± 4.2 vs 9.6 ± 3.4°, p = 0.011, **FIGURE 2b**).

**FIGURE 1.**
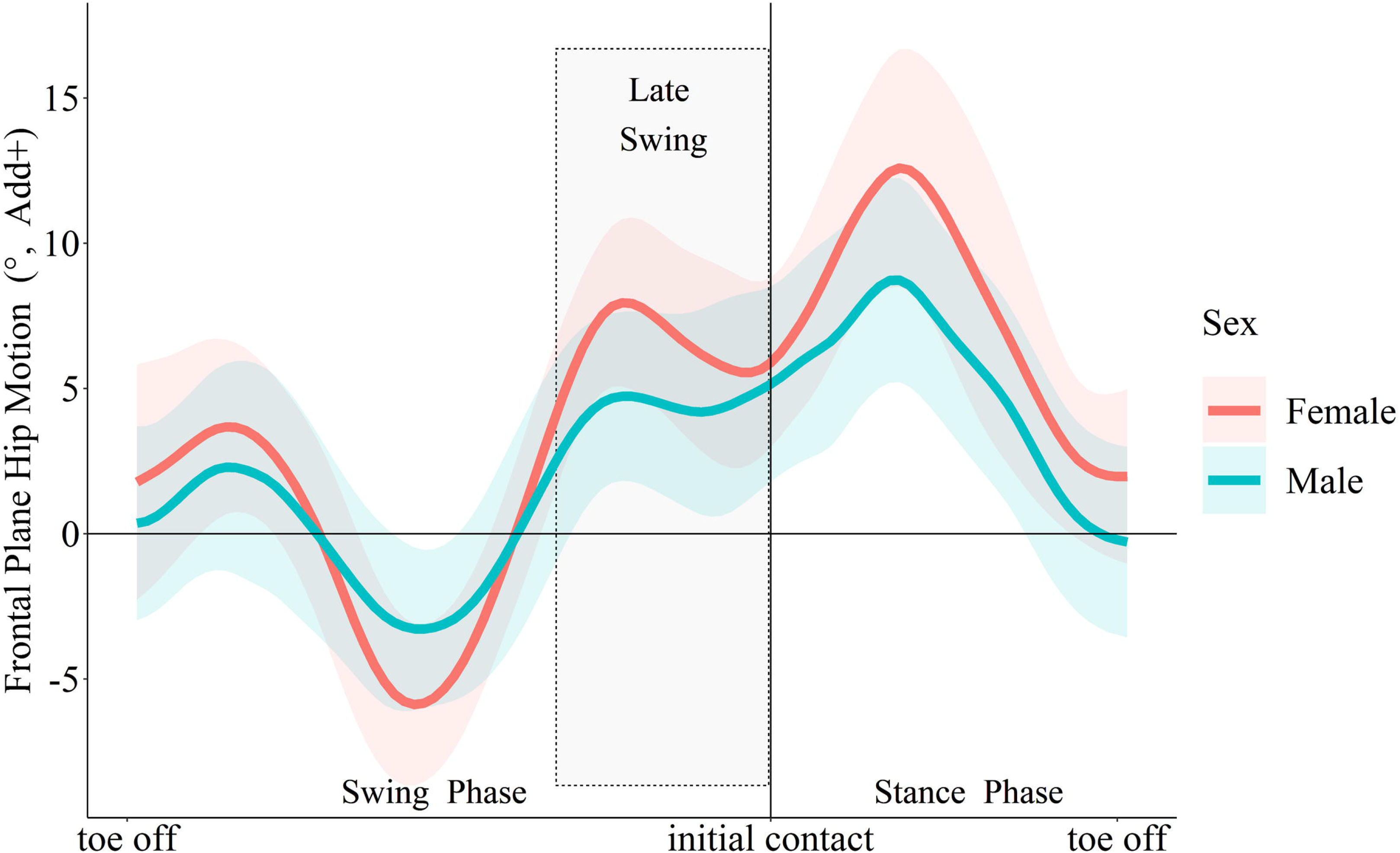
Sex difference in frontal plane hip motion during running. Data are mean ± SD.

**FIGURE 2.**
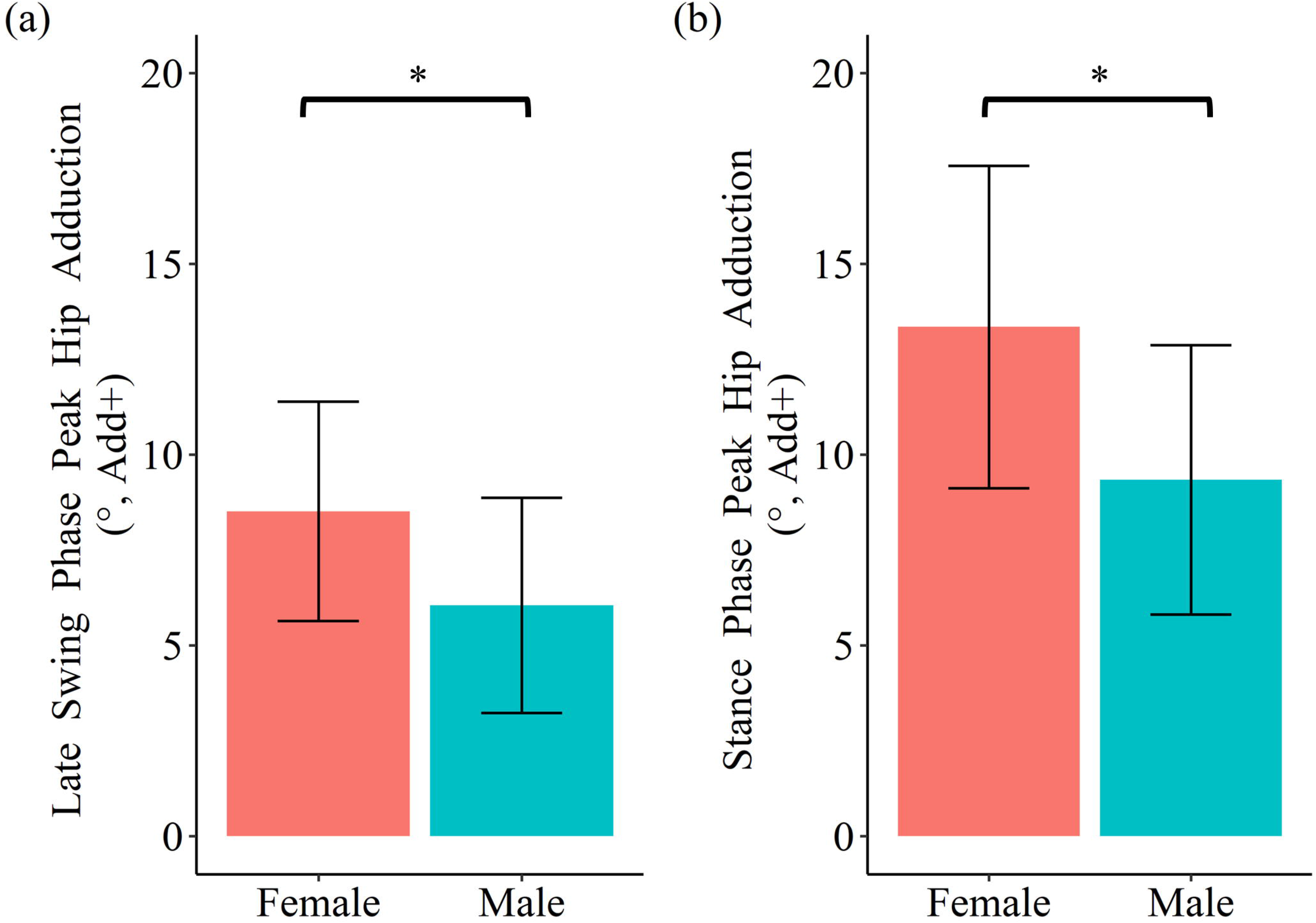
Sex difference in peak hip adduction angles during late swing phase (a) and stance phase (b) of running. Data are mean ± SD. *: *p*<0.05

Results of the multiple regression analysis did not reveal a significant sex interaction (p>0.05). As such, data from males and females were pooled in the final model. Peak hip adduction during late swing was found to predict peak hip adduction during stance (r = 0.63, p < 0.001, **FIGURE 3**).

**FIGURE 3.**
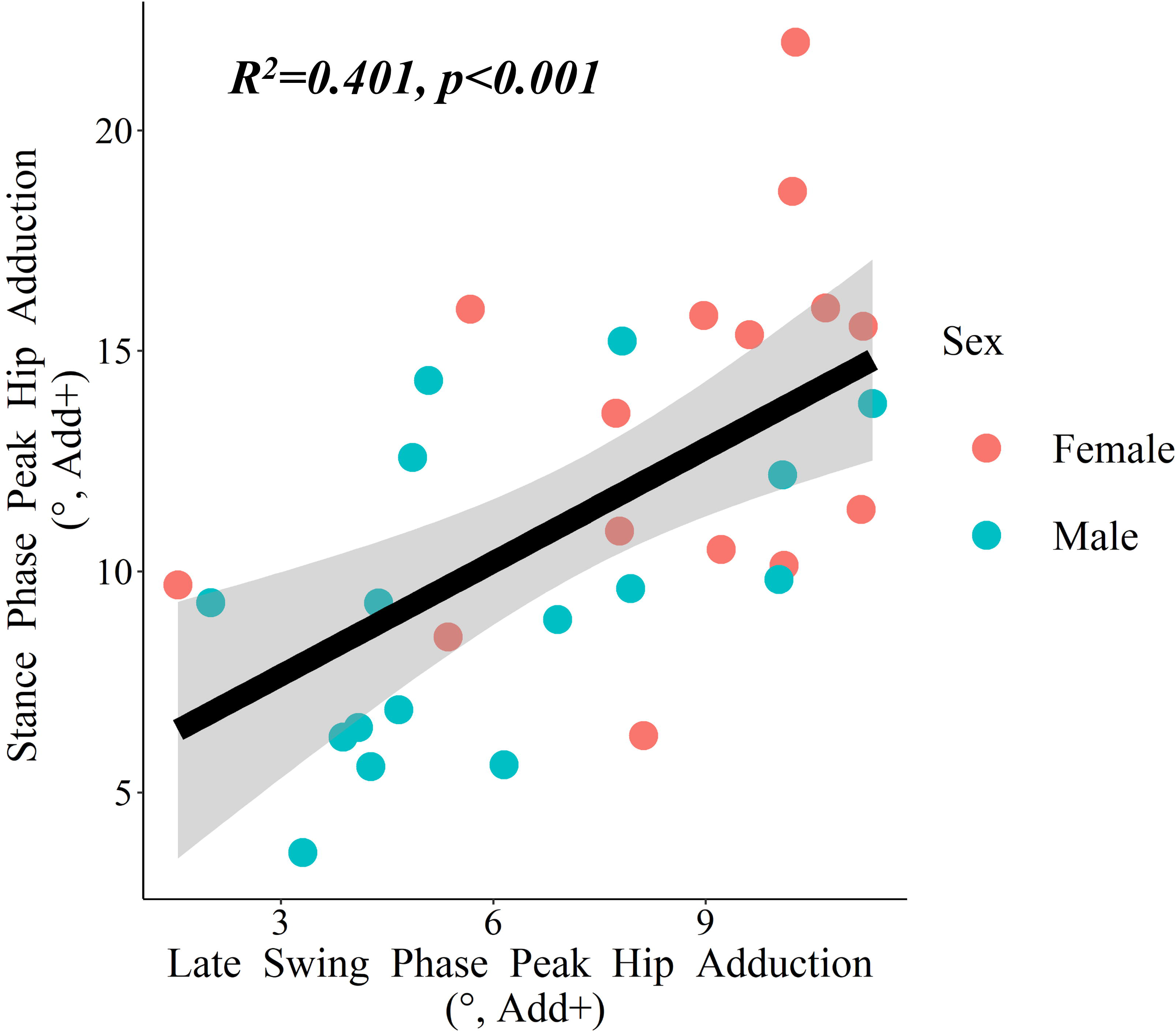
Correlation of late swing phase peak hip adduction to stance phase peak angles during running. Pink dots represent females and blue dots represent males. Grey shading represents the standard error of the slope.

## DISCUSSION

The purpose of the current study was to evaluate sex differences in peak hip adduction during the late swing and stance phases of running and determine whether peak hip adduction during late swing was predictive of peak hip adduction during stance. Consistent with our hypotheses, significant sex differences in peak hip adduction were found during both stance and late swing phases. In addition, peak adduction during late swing was found to be predictive of peak hip adduction during stance.

Our finding of sex differences in peak hip adduction during the stance phase of running is consistent with previous studies (7,11,14–16,28). On average, the females in the current study exhibited approximately 4 degrees greater peak hip adduction during stance than males. This difference is similar in magnitude to previous studies that have reported sex differences peak hip adduction ranging from 4 to 6 degrees (7,11,14–16,28). Although small, differences in hip adduction in this range are considered clinically meaningful as prospective studies involving female runners have reported that peak hip adduction differences of 4-5 degrees can discriminate among runners who developed iliotibial band syndrome (6) and patellofemoral pain (2) from those who do not.

Regarding peak hip adduction during late swing, females exhibited 2 degrees greater hip adduction on average compared to males. Although statistically significant, this finding is smaller in magnitude to previously reported sex differences in hip adduction at initial contact, which have averaged about 4 degrees (7,13,16). Differences among studies may be due to the fact that the current investigation evaluated peak hip adduction during the last one-third of the swing phase as opposed to initial contact. However, our results related to late swing phase hip adduction are in agreement with graphical results of Chumanov et al. (28) and Schache et al. (29).

The novel finding of the current study was that peak hip adduction during late swing was predictive of peak hip adduction during stance. Specifically, 40% of the variance in stance phase peak hip adduction could be explained by late swing values. Interestingly this percent of explained variance for peak hip adduction during the stance phase of running is considerably higher than what has been reported for hip abductor strength. For example, studies that have reported significant associations between hip abductor strength and hip adduction during running have reported R^2^ values up to 16% (12,21–24). Taken together, our results suggest that sex difference in hip adduction during the stance phase of running has its origin, at least in part, during late swing.

Although determining reasons for increased hip adduction during late swing of running was beyond the scope of the current study, it is interesting to speculate on potential causes. One potential contributing factor may be related to known sex differences in pelvis morphology. For example anatomical differences such as pelvis width, bilateral hip joint center distance, acetabular orientation, and femoral neck-shaft angle could result in the need for greater hip adduction during late swing to adequately position the foot relative to the center of mass to achieve stance phase stability. Although previous studies have evaluated the association between measures of pelvis and femur morphology with hip adduction during the stance phase of running (e.g. femoral neck shaft angle (24), hip joint center width (28), bi-greater trochanteric distance (30)), the reported amount of variance in hip adduction explained by these morphologic measures have been non-significant (24,28) or very low (R^2^=0.05) (30). It is possible that pelvis and femur morphology may be more predictive of hip adduction during swing, as motion during this phase is less likely to be influenced by additional variables such as external demand (i.e., adduction moment) and hip abductor strength. Several limitations need to be considered when interpreting the results of this study. First, the participants from our study were young, healthy recreational runners. Caution is needed when generalizing results of this study to injured runners or older populations. In addition, we only examined heel strike runners. As such, our findings may not apply to forefoot or midfoot strike runners. Lastly participants were instructed to run at 4 m/s which is a moderate speed. Whether or not our results would be relevant to faster speed running or sprinting remains to be determined.

## SUMMARY

When compared to males, females demonstrated greater amounts of hip adduction during the stance and swing phases of running. In addition, peak hip adduction in late swing was predictive of respective peak angles during the stance phase of running. As such, the hip position before ground contact appears to be an important determinant of stance phase hip adduction. Future studies should consider potential causes of these kinematics during late swing (e.g. bony morphology, muscle activation) to better understand running injuries associated with excessive stance phase hip adduction. In addition, future work should be directed towards understanding if hip re-positioning during late swing can be used as a potential method of effecting peak hip adduction during stance.

## ACKNOWLEDGEMENTS

This study was funded by American College of Sports Medicine (ACSM), International Society of Biomechanics, American Society of Biomechanics, and University Voucher Program Grants.

There is no conflict of interest to disclose. The results of the present study do not constitute endorsement by ACSM. The results of the study are presented clearly, honestly, and without fabrication, falsification, or inappropriate data manipulation.

